# Female ornamentation and the fecundity trade-off in a sex-role reversed pipefish

**DOI:** 10.1101/290387

**Authors:** Kenyon B. Mobley, John R. Morrongiello, Matthew Warr, Diane Bray, Bob B. M. Wong

## Abstract

Sexual ornaments found only in females are a rare occurrence in nature. One explanation for this is that female ornaments are costly to produce and maintain and, therefore, females must trade-off resources related to reproduction to promote ornament expression. Here, we investigate whether a trade-off exists between female ornamentation and fecundity in the sex-role reversed, wide-bodied pipefish, *Stigmatopora nigra*. We measured two components of the disk-shaped, ventral-striped female ornament, body width and stripe thickness. After controlling for the influence of body size, we found no evidence of a cost of belly width or stripe thickness on female fecundity. Rather, females that have larger ornaments have higher fecundity and thus accurately advertise their reproductive value to males without incurring a cost to fecundity. We also investigated the relationship between female body size and egg size and found that larger females suffered a slight decrease in egg size and fecundity, although this decrease was independent of female ornamentation. More broadly, considered in light of similar findings in other taxa, lack of an apparent fecundity cost of ornamentation in female pipefish underscores the need to revisit theoretical assumptions concerning the evolution of female ornamentation.

## Introduction

In many species, males are the more competitive sex and are adorned with elaborate ornaments that are used as visual signals to attract potential mates (Andersson 1994). Male ornaments are thought to arise through sexual selection via female mate choice with more ornamented males favored by choosy females (Andersson 1994; Hill 2014). Such preferences are generally ascribed to the high fitness costs of bearing such ornaments (Zahavi 1975, 1977; Grafen 1990; Walther & Clayton 2005; Hill 2014). Specifically, it has traditionally been assumed that only high quality individuals should be able to bear the cost of maintaining the most elaborate ornaments, causing the degree of ornament elaboration to function as an honest indicator of an individual’s quality (Zahavi 1977; Kodric-Brown & Brown 1984; Nur & Hasson 1984; Grafen 1990; Walther & Clayton 2005). However, according to more recent theoretical models, costs alone may not be sufficient to explain the persistence of exaggerated ornaments in nature (Tazzyman *et al*. 2013, 2014).

In contrast to what is known about male sexual ornaments, far less is known about the evolution of female ornaments (Clutton-Brock 2007, 2009). In some rare instances, females have evolved elaborate ornamentation without a comparable ornament in males. Female-specific ornaments have evolved in diverse taxa, such as insects (LeBas *et al*. 2003; Bussiere *et al*. 2008; Takahashi & Watanabe 2011; Wheeler *et al*. 2012; Hopkins *et al*. 2015), crabs (Baldwin & Johnsen 2012), fishes (Amundsen & Forsgren 2001; Rosenqvist & Berglund 2011), reptiles (LeBas & Marshall 2000; Weiss 2006), birds (Amundsen 2000b; Roulin *et al*. 2003; Gladbach *et al*. 2010), and mammals (Huchard *et al*. 2009). Although it is generally believed that female-specific ornamentation evolves similarly to male ornamentation through the process of mate choice and sexual selection (Amundsen 2000a, b; Clutton-Brock 2007, 2009), females putatively evolve ornaments via social interactions including signaling dominance or social selection (i.e., competition for resources, including mates) (West-Eberhard 1979; LeBas 2006; Lyon & Montgomerie 2012; Tobias *et al*. 2012).

One explanation for its rarity is that female ornamentation is costly, particularly with respect to fecundity (Fitzpatrick *et al*. 1995; Chenoweth *et al*. 2006; Clutton-Brock 2007, 2009). In general, female investment in reproduction is greater than in males, and female quality is based on her fecundity, the quality of her eggs, and/or parental care investment (Trivers 1972; Clutton-Brock 2009). Female ornaments are therefore not favored to evolve if their production is costly in terms of future investment into offspring (Fitzpatrick *et al*. 1995). Due to a general positive body size-fecundity relationship in many species (e.g., Barneche *et al*. 2018), larger females may represent a higher reproductive value to prospective mates and perhaps additional ornaments may be unnecessary (Hopkins *et al*. 2015). However, additional ornaments may serve to amplify information about her quality to choosy males, particularly if ornaments accentuate her body size and, hence, her fecundity (Rosenqvist & Berglund 2011). Female ornaments may also be used as a signal in female-female competition where females compete for high quality males that provide direct and indirect genetic benefits to offspring (Bernet *et al*. 1998; Rosvall 2011).

If ornament expression is an accurate indicator of fecundity, then the rate of increase of the relationship between ornament expression and fecundity should be equal (Simmons & Emlen 2008). Deviations from a positive linear relationship should indicate a trade-off between ornament expression and fecundity. If costs increase with ornament expression, individuals displaying large ornaments would suffer reduced fecundity. Alternatively, if costs decrease with ornament expression such that smaller individuals are saddled with a higher cost to producing the ornament, or if producing larger ornaments are cheaper in larger individuals, we would expect to see proportionately larger ornaments in larger individuals.

To date, only a handful of studies have investigated the ornament-fecundity trade-off in species that possess female ornaments, and evidence for such a trade-off is generally lacking. For example, a study on the dance fly, *Rhamphomyia tarsata*, found a linear relationship in fecundity and the length of pinnate scales with female body size (LeBas *et al*. 2003). Similarly, studies on female ornamentation in the striped plateau lizard, *Sceloporus virgatus*, have revealed a positive relationship between ornaments, fecundity and offspring quality (Weiss 2006; Weiss *et al*. 2009). In horned beetles, *Onthophagus sagittarius*, female body size was the most reliable predictor of maternal quality, yet developing relatively large horns did not impart a cost to fecundity (Simmons & Emlen 2008). In the upland goose, *Chloephaga picta leucoptera*, female-specific coloration was related to clutch and egg volumes (Gladbach *et al*. 2010), while in glow worms, *Lampyris noctiluca*, the intensity of the glow emitted by females was positively associated with body size and fecundity (Hopkins *et al*. 2015). In at least one case, the eider duck, *Somateria mollissima*, female-specific plumage was unrelated to clutch size or the phenotypic quality of females (Lehikoinen *et al*. 2010). Thus, despite the theoretical costs of female ornamentation on fecundity, little empirical evidence exists to support such a scenario.

Members of the family Syngnathidae (pipefish, seahorses, and seadragons) offer an outstanding opportunity to investigate the evolution of female ornamentation because a remarkable diversity of female ornaments has evolved in several lineages, ranging from temporary courtship ornaments to extreme sexual dimorphism, brilliant permanent markings and flashy displays (Dawson 1985; Kuiter 2009; Rosenqvist & Berglund 2011). Female ornaments ostensibly evolved in syngnathids because of the unique reproductive mode of this group: male pregnancy. Males provide all parental care and species with enclosed brood pouches provide protection, osmoregulation and nutrition to developing embryos (Haresign & Shumway 1981; Ripley & Foran 2006; Partridge *et al*. 2007; Ripley & Foran 2009; Kvarnemo *et al*. 2011). In most syngnathid species, male pregnancy decreases the rate at which males can mate but not females, thereby increasing competition between females for access to mating opportunities (Berglund *et al*. 1986a; Kvarnemo & Ahnesjö 1996). Female mate-limitation creates a biased operational sex ratio and results in sex-role reversal where sexual selection acts more strongly on females than on males (Berglund *et al*. 1986b; Vincent *et al*. 1992; Jones *et al*. 2000). Consistent with sexual selection theory, the genetic mating system across syngnathid species predicts the strength of selection for exaggerated ornamentation, i.e., females from highly polyandrous species have the most striking ornaments (Jones & Avise 2001; Rosenqvist & Berglund 2011).

This study aimed to investigate the trade-off between female ornamentation and fecundity in the wide body pipefish, *Stigmatopora nigra*, Kaup 1856.

This species is ideal because females possess an exaggerated ornament that females display to males during courtship. The ornament is a large dorsoventerally flattened, disc-shaped belly with alternating light and dark ventral stripes (Fig. 1). The width of the ornament and stripes can be measured directly, and fecundity can be obtained by counting mature ova in the ovaries of females. Accordingly, we explored the relationship between female fecundity and the size and stripe pattern of the female ornament. We also investigated the relationship between mean egg size and ornament expression. We predicted that if there is no fecundity cost to ornament expression, then a positive linear relationship between ornament expression and fecundity or egg size should be observed.

**Figure 1.**
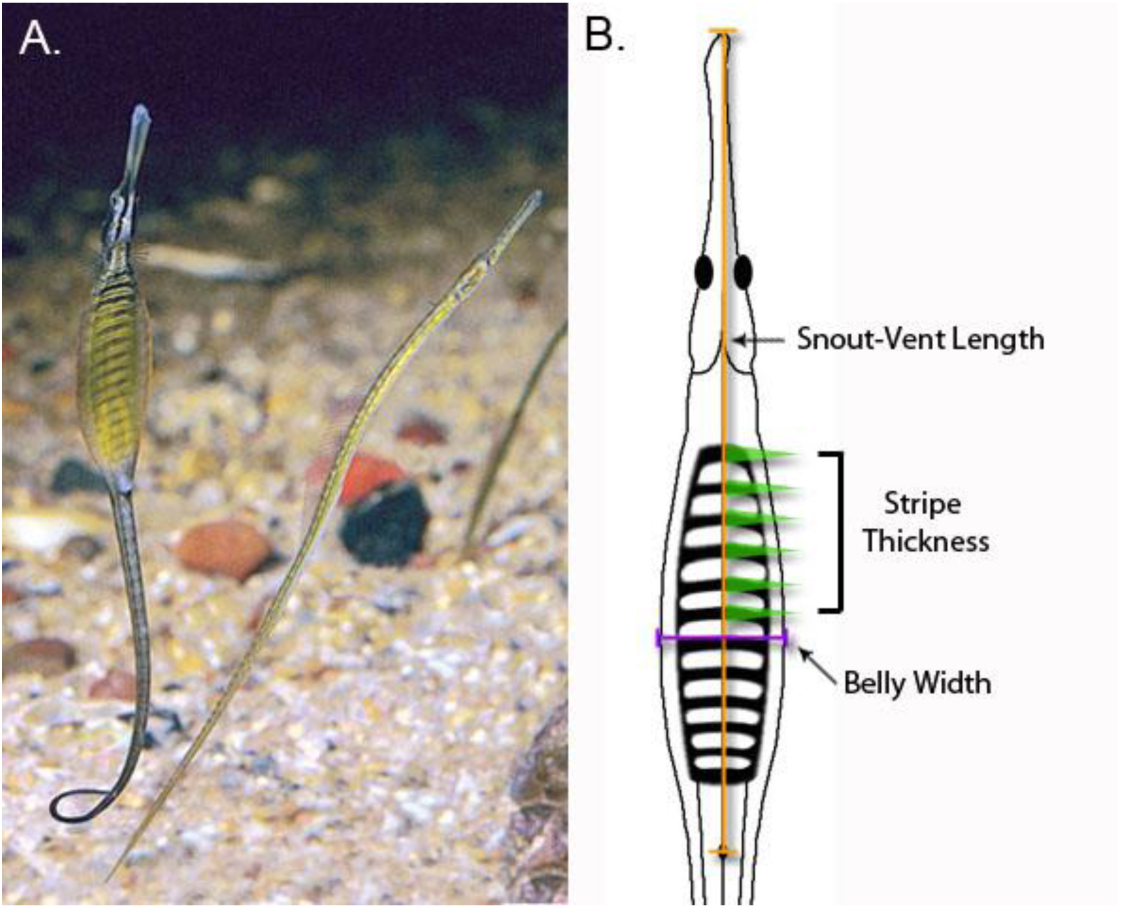
A. Female (left) and male (right) *Stigmatopora nigra*. The female is displaying her striped belly ornament to the male. B. Schematic diagram of measurements used in this study. Snout-vent length is estimated from ventral photographs from the tip of the rostrum to the anal pore. Stripe thickness is the mean width of the first six dark stripes. Belly width is calculated as the widest part of the body. Photography © Rudie Kuiter, used with permission.

## Material and methods

### Study species

The wide body pipefish, *Stigmatopora nigra* (Kaup, 1856), occurs in bays, estuaries and shallow coastal waters of southern Australia and New Zealand (Dawson 1985). Wide body pipefish breed throughout the year in shallow seagrass beds and their abundance and the proportion of pregnant males reach their peak in September–January (Duque-Portugal 1989). Females possess a wide, laterally compressed body, a darkly pigmented dorsum and a striped, ventral ornament that is displayed to males during courtship (Fig. 1a). Occasionally, females have an additional fleshy fold on the lateral edges of the ornament (Dawson 1985). Males do not possess a wide body and lack stripes on their brood pouch (Fig. 1a). Males have a semi-inverted pouch enclosure and care for offspring until parturition (Dawson 1985; Wilson *et al*. 2003). The mating system of the species is unknown due to lack of molecular parentage studies conducted on this species. However, because of the strong sexual dimorphism and possession of a female ornament, the species is most likely polyandrous where males mate with a single female while females can mate with multiple males (Jones & Avise 2001). This species is also putatively sex-role reversed with respect to sexual selection (i.e., sexual selection acting more strongly on females than males) similar to other pipefish species that display female ornaments (Berglund *et al*. 1986b; Jones *et al*. 2000).

### Sample collections

Adult *S. nigra* used in the study were museum specimens collected either by using drop or seine nets at Grassy Point in Port Phillip Bay on the Bellarine Peninsula, Victoria, Australia (38°07’ S, 144°41’ E). Specimens were sampled across multiple years (1997, 1999, 2005, and 2006) during September – January as part of a series of unrelated studies (Jenkins & Hamer 2001; Jenkins *et al*. 2002; MacReadie *et al*. 2009). These fish were euthanized immediately after capture using either a 99% ethanol solution or benzocaine before being preserved in 70% ethanol and stored in the collection at Museum Victoria.

### Morphological measurements

Female pipefishes were photographed using a Nikon D80 digital SLR camera for morphological measurements. Females were placed on a foam board covered by a sheet of laminated paper with 1mm grid lines for scale. The females were then pinned down flat with their ventral side exposed, as this allowed for accurate measures of both body size and ornamental traits. Measurements were then taken from the photographs using the image analysis software ImageJ (http://rsb.info.nih.gov/ij/). Because 19% of females and 24% of males had broken tails, we used snout-vent length (SVL, tip of rostrum to anal pore, Fig. 1b) as a measure of body size as opposed to total length (TL, tip of rostrum to tip of tail). Snout-vent length was highly correlated to total length in both sexes (female: r = 0.944, t_1,102_ = 28.90, p < 0.0001; male: r = 0.938, t_1,54_ = 19.88, p < 0.0001). We obtained two different measures of female ornamentation: belly width and stripe thickness. Belly width was measured from the widest point perpendicular to SVL to ensure uniformity of measurements (Fig. 1b). We also measured belly width in males. For stripe thickness, the width of dark stripes at the midpoint of the SVL axis was measured (Fig. 1b). While preservation caused fading of the dark stripes on many females, the first six stripes were visible for the majority of the females. Therefore, the mean thickness of the first six stripes was used as the measurement for stripe thickness.

### Dissections and egg size measures

After photography, females were placed in a petri dish and submerged in water to prevent desiccation. Ovaries were dissected from females and eggs gently separated from ovarian tissue using tweezers, enumerated to estimate fecundity, and the egg diameters measured under 40X magnification using a 0.1mm graticule. Due to the ovoid shape of most eggs, two perpendicular measures of diameter were taken and averaged in order to estimate mean egg size.

Female ovaries contained both immature and mature eggs, although preservation made it difficult to identify the two. Counting immature eggs would overestimate a female’s current fecundity, or potential clutch size, and thus eggs available to a male during mating. Therefore, we developed a method to estimate which eggs were mature in female ovaries, and hence her fecundity using the size range of newly laid eggs (without eyespots) located within the brood pouch of ethanol-preserved males. Eggs from each male’s brood pouch were dissected, enumerated for an estimate of male reproductive success and the diameters measured under 40X magnification using a 0.1mm graticule. We assumed that each male has eggs from just one female as developing embryos within male brood pouches were uniformly distributed throughout the pouch and at similar developmental stages. We then used linear regression to find the relationship, across males, between the minimum and maximum egg sizes in a pouch. A significant positive relationship was found between the minimum and maximum egg size found in a male’s brood pouch (r = 0.518, t_1,57_ = 4.57, p < 0.0001). The maximum egg diameter was found to correlate to 0.613*the minimum egg diameter found in a male’s pouch + 0.587. We applied this equation to predict the smallest mature egg size of a female, dependent on the largest egg size found in her ovary. We calculated female fecundity as the sum of all eggs within a female’s ovaries that were considered mature using this method.

### Statistical analysis

We analysed data from 104 females and 59 males for which all metrics were available (SVL, fecundity, egg size, belly width and stripe thickness in females; egg number and egg size in males, Table 1). Basic statistics were calculated in R (R Core Team 2017). Variables were tested for normality and equal variances (Levene) and transformed, if necessary, to satisfy assumptions. Means are reported ± standard error of the mean.

**Table 1.** The number (n) of male and female *Stigmatopora nigra*, mean snout-vent length (SVL), mean number of mature eggs in female ovaries (i.e., fecundity), mean number of eggs in male brood pouch, mean egg size, mean belly width and mean stripe thickness of females. All means are reported ± one standard error of the mean.

|  | n | SVL (cm) | Number of eggs | Egg size (mm) | Belly Width (mm) | Stripe Thickness (mm) |
| --- | --- | --- | --- | --- | --- | --- |
| Female | 104 | 4.47 ± 0.06 | 32.9 ± 1.4 | 0.92 ± 0.01 | 4.53 ± 0.10 | 0.76 ± 0.01 |
| Male | 59 | 3.61 ± 0.08 | 32.7 ± 1.9 | 0.95 ± 0.01 | 2.15 ± 0.05 | -- |

Based on SVL measurements of sexually mature adults, we found a unimodal normal distribution of male (Shapiro-Wilk W test: W = 0.9782, p = 0.3672) and female body sizes (Shapiro-Wilk W test: W = 0.9839, p = 0.2397) suggesting that *S. nigra* only live for one year. A unimodal distribution is corroborated with yearly sampling of this species (Duque-Portugal 1989) and similar to other temperate pipefishes (Mobley *et al*. 2010; Braga Goncalves *et al*. 2017). Larger females had proportionately larger ornaments (SVL vs. belly width: r = 0.824, t_1,102_ = 14.69, p < 0.0001; SVL vs. mean stripe thickness: r = 0.768, t_1,102_ = 12.11, p < 0.0001). Therefore, ornament traits were standardized for female body sizes by using the raw residuals from ordinary least squares regressions of ornament ~ SVL.

To investigate the relationship between female ornaments and fitness correlates, we modelled female fecundity and egg size as a function of female size-adjusted ornamentation and SVL using linear mixed effects models fitted in the lme4 package V 1.1-17 in R (Bates *et al*. 2015). We included a random intercept for sample year to account for potential among-year differences in reproductive investment driven by unmeasured environmental conditions. We compared a series of increasingly complex models (fitted with maximum likelihood) that included linear and quadratic terms for ornamentation and SVL. We then used this modelling framework to explore the relationship between fecundity and egg size, accounting for SVL. We used Akaike Information Criterion corrected for small sample size (AICc, Burnham & Anderson 2002) to select the best fit model. Models with ΔAICc < 2 were considered to have similar support. Fecundity data was natural log-transformed to satisfy model assumptions and predictor variables were mean-centered to facilitate interpretation of polynomial terms. Parameter estimates and 95% credible intervals were derived from the posterior distribution of the fixed effects in the best models (fitted with restricted maximum likelihood, REML) using 1000 model simulations generated by the arm package for R.

## Results

Sexual dimorphism is apparent in this species: females were larger and have bellies that were twice the width of males, on average (Analysis of variance (ANOVA) SVL: F_1,161_ = 82.6, p < 0.0001; ANOVA belly width: F_1,161_ = 440.8, P < 0.0001; Table 1). Females varied widely in fecundity (14-89 eggs), belly width (2.2-6.7mm), and stripe thickness (0.38-1.1mm). There was a significant positive relationship between the two ornamental traits, belly width and stripe thickness, when accounting for body size (Analysis of covariance (ANCOVA) stripe thickness: F1,101 = 32.39, P < 0.0001; SVL: F1,101 = 40.79; P < 0.0001, no interaction between stripe thickness and SVL). Male brood pouches contained between 1 and 76 eggs (mean number of eggs: 32.7, Table 1).

The best fecundity model predicted by morphological traits included linear and quadratic terms for standardized belly width and a linear term for SVL (Table 2). A model with similar support (ΔAICc=0.7) included just linear terms for standardized belly width and SVL. Larger females were more fecund (β(SVL)= 0.297 [0.195 to 0.395 95%CI], Fig. 2a). Females with relatively larger belly widths also had higher fecundity (β(stand.BW)= 2.602 [1.703 to 3.617 95%CI], β(stand.BW2)= −8.518 [−18.266 to 1.158 95%CI], Fig. 2b). There was little support for the positive linear relationship between fecundity and standardized stripe thickness (ΔAICc to best Fecundity model=34.1; Fig. 2c). The best egg size model included linear and quadratic terms for SVL (Table 2). Medium sized females had the largest eggs (β(SVL)= 1.066 [0.597 to 1.498 95%CI], β(SVL2)= −0.116 [−0.165 to −0.063 5%CI], Fig. 2d). There was no relationship between standardized belly width or stripe thickness and egg size (Fig. 2e, 2f).

**Table 2.**
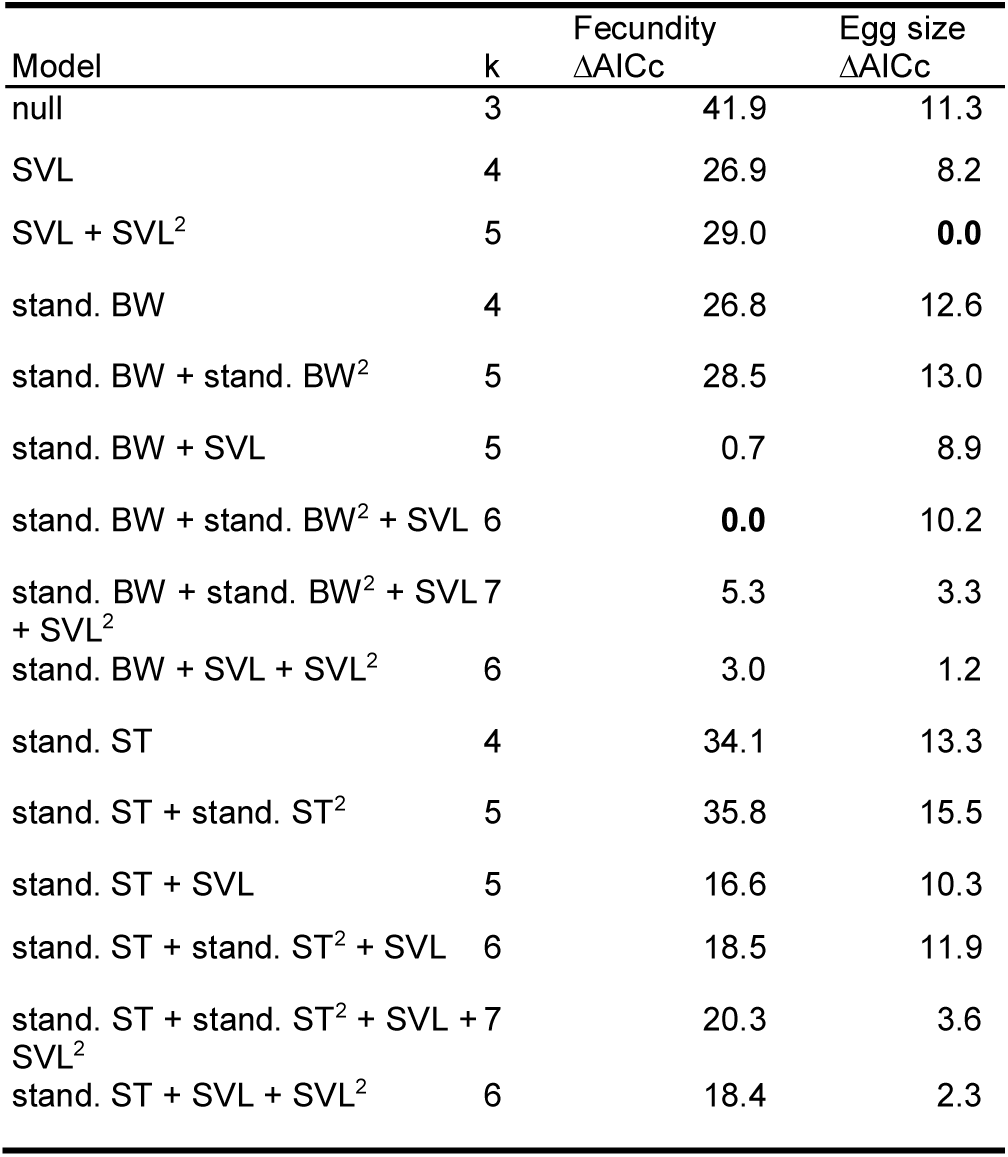
Results of AlCc based model selection for female fecundity and mean egg size. SVL=snout-vent length, stand.BW=standardised belly width; stand.ST=standardised stripe thickness; k=number of model parameters. The best model for each reproductive measure (ΔAICc= 0) is highlighted in bold.

**Figure 2.**
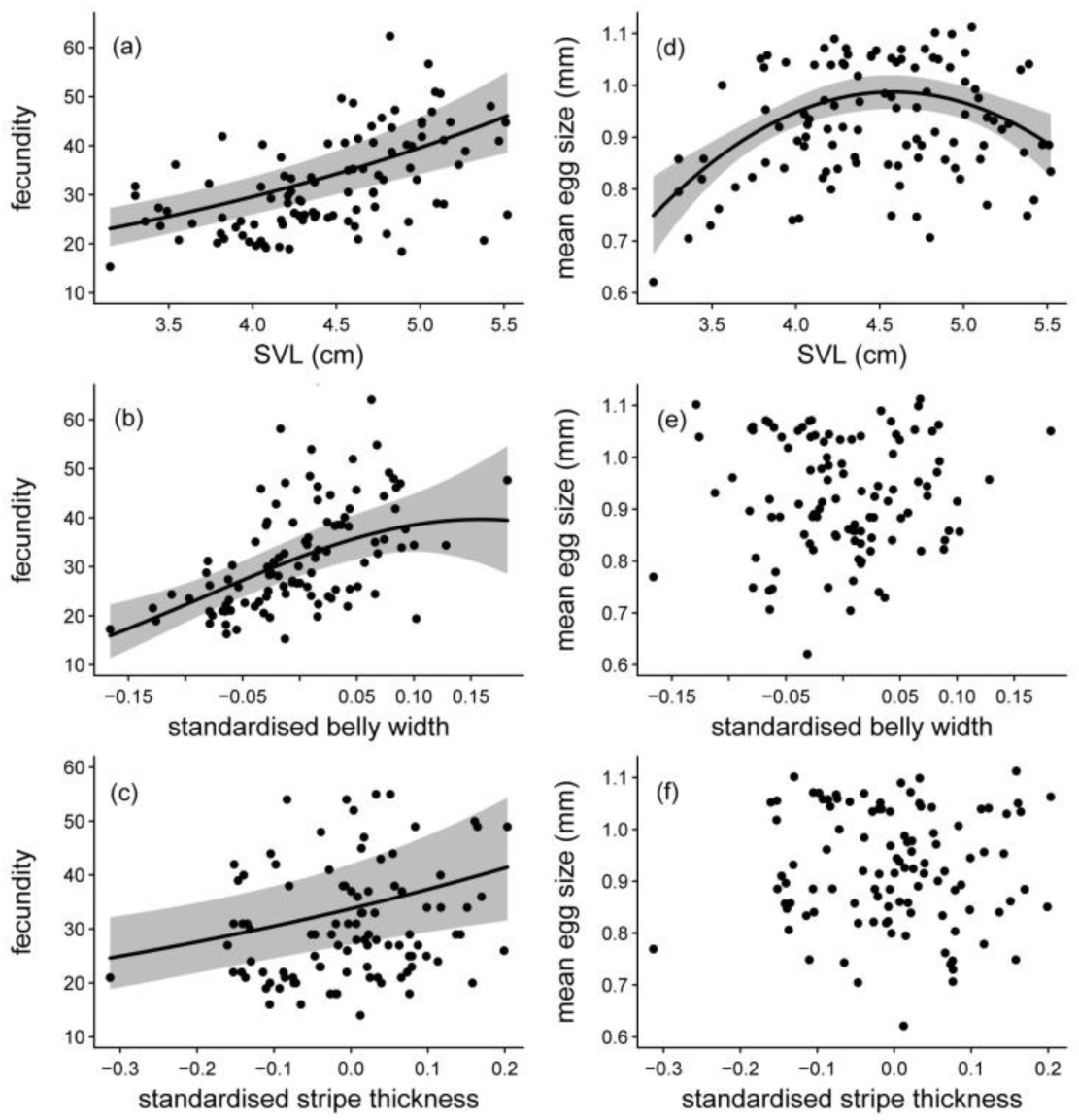
Predicted relationship (grey areas are ± 95% CI) between (a) fecundity and snout-vent length (SVL); (b) fecundity and standardized belly width (see methods); (c) fecundity and standardized mean stripe thickness (see methods); (d) mean egg size and SVL; (e) mean egg size and standardized belly width; (f) mean egg size and standardized stripe thickness. Points in (a) and (b) represent partial effects from multiple mixed model regression (other covariates are held at mean values). Points in c-f are observations.

The best fecundity-egg size model included a linear term for SVL and a quadratic term for egg size (Table 3). After accounting for larger females having higher fecundity, we found that fecundity was curve-linearly related to mean egg size (β_(egg)_= 0.972 [0.436 to 1.490 95%CI], 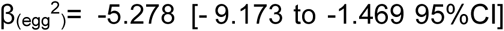. Females of intermediate fecundity had the largest eggs (Fig. 3).

**Table 3.** Results of AICc based model selection for female fecundity and egg size relationship. SVL= snout-vent length; k= number of model parameters. The best model (ΔAICc= 0) is highlighted in bold.

| Model | k | $\Delta\text{AICc}$ |
| --- | --- | --- |
| null | 3 | 32.1 |
| SVL | 4 | 17.1 |
| SVL + SVL<sup>2</sup> | 5 | 19.2 |
| Egg size | 4 | 14.3 |
| Egg size + Egg size<sup>2</sup> | 5 | 6.6 |
| SVL + Egg size | 5 | 4.6 |
| SVL + Egg size + Egg size<sup>2</sup> | 6 | **0** |
| SVL + SVL<sup>2</sup> + Egg size | 6 | 6.6 |
| SVL + SVL<sup>2</sup> + Egg size + Egg size<sup>2</sup> | 7 | 1.2 |

**Figure 3.**
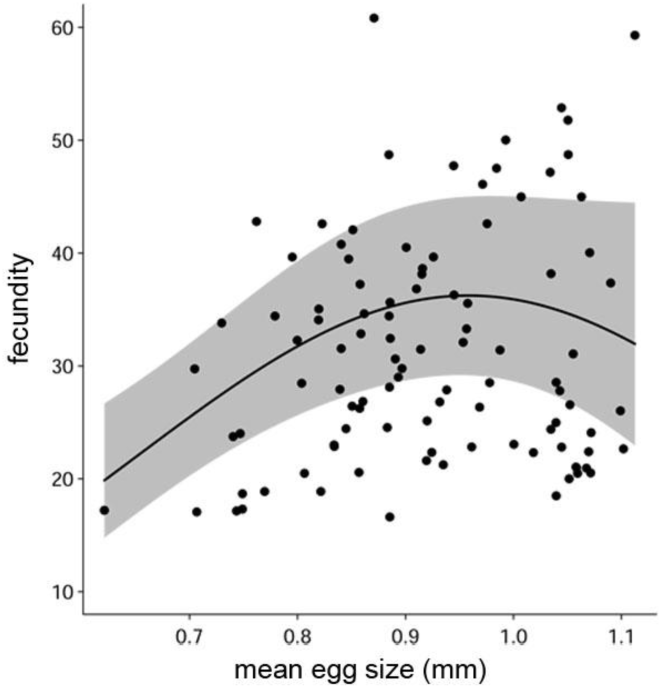
Predicted relationship (grey areas are ± 95% CI) between fecundity and mean egg size, accounting for SVL. Points represent partial effects from multiple mixed model regression (SVL held at its mean value).

## Discussion

In this study we explored the potential relationship between female ornament expression and fecundity in a species of pipefish where females possess a highly exaggerated belly ornament not found in males. We uncovered strong evidence for a positive linear relationship between ornament expression and fecundity, with larger females having proportionately larger ornaments and higher fecundity than their smaller counterparts. Ornaments often scale by traits such as body mass and/or size and are often condition dependent (Johnsen *et al*. 1996; Amundsen *et al*. 1997; Kotiaho 2000; Simmons & Emlen 2008). In such cases, larger or better condition females should have proportionately larger ornaments. Based on our findings, we can conclude fecundity costs to ornament expression in S. *nigra* is negligible. Instead, body size, as well as belly width and stripe thickness accurately reflect female fecundity. These results add to a growing body of literature that have similarly found no evidence of fecundity costs in species with female-specific ornamentation (LeBas *et al*. 2003; Weiss 2006; Simmons & Emlen 2008; Weiss *et al*. 2009; Gladbach *et al*. 2010; Lehikoinen *et al*. 2010; Hopkins *et al*. 2015). Together, they challenge traditional assumptions about fecundity costs as an explanation for the scarcity of female ornament expression.

We also investigated the relationship between female body size and egg size and found that, independent of female ornamentation, larger females tended to produce smaller eggs than average size females. Such a relationship may allow larger females to produce more eggs than smaller females, but, in so doing, may also result in fitness costs to offspring. Indeed, egg size is often related to offspring fitness in a wide range of taxa, with larger eggs bestowing higher fitness benefits to developing young (Green 2008; Weiss *et al*. 2009; Krist 2011). This also appears to be the case in pipefishes (Ahnesjö 1992), although females can also strategically mate or distribute resources to eggs and clutches depending on male quality (Braga Gonçalves *et al*. 2010; Paczolt & Jones 2010; Mobley *et al*. 2011). One potential explanation for the egg size relationships observed in our study is that large female *S. nigra* may have fewer resources to provision future broods and, hence, egg size may decrease as a result of resource depletion (Wiklund & Karlsson 1984). However it is currently unknown if *S. nigra* are capital breeders (i.e., pay for reproduction based on stored resources, *sensu* (Stephens *et al*. 2009)) or if larger females are more successful at breeding. Alternatively, larger females may strategically decrease their egg size to increase fecundity (Smith & Fretwell 1974; Parker & Begon 1986). Reduction in egg and/or clutch size is hypothesized to be adaptive if the cost of reproduction declines with increasing age and if age-selective mortality is low relative to reproduction-dependent mortality (Begon & Parker 1986). Finally, it is worth noting that all females were collected in the wild and may have recently mated, potentially affecting the fecundity and egg size of particular females. Currently, the ovarian type in *S. nigra* is unknown, although at least two types have thus far been described in pipefishes: the so-called asynchronous type, where females produce small numbers of eggs continuously (e.g. *Syngnathus scovelli* and *Syngnathus typhle:* (Begovac & Wallace 1987)), and the group-synchronous type (e.g. *Nerophis ophidion:* (Sogabe & Ahnesjö 2011); *Corythoichthys haematopterus:* (Sogabe *et al*. 2008)), where ovaries are mature in distinct clutches. Because mature egg size was deduced on the relative size of eggs within the ovaries based on the size of eggs within males, it is possible that we overestimated fecundity by including some non-mature eggs. Thus, if larger females recently mated and have a high proportion of non-mature eggs in the ovaries, this may account for the reduced mean egg size in the largest females.

If possessing female-specific ornaments isn’t costly in terms of future offspring production, this then begs the question, why do we not see female ornaments evolve in more systems? High costs to fecundity from ornamentation may still exist in species that do not have appreciable female-specific ornamentation, thus precluding the evolution of such structures in the first place. Moreover, if body size is the best predictor of fecundity, then additional ornaments may be superfluous (Hopkins *et al*. 2015), especially if they are also costly in other ways (e.g. increased risk of predation). Among species that do possess female-specific ornaments, increased signal efficacy and/or condition dependence may help to explain why female ornaments are maintained when no clear cost to fecundity is apparent. For example, larger individuals may be more effective at advertising and, as a result, signal strength may increase with body size (Tazzyman *et al*. 2013, 2014). Alternatively, if condition (i.e. fecundity) increases with body size, then we would expect to see larger females honestly advertising their condition (Tazzyman *et al*. 2014). It is interesting to note that the lateral stripe pattern of S. *nigra’s* ornament is common in pipefishes and may accentuate body depth putatively via the so-called Helmholtz illusion (Berglund 2000; Rosenqvist & Berglund 2011). This optical striped illusion creates the appearance that the belly is wider than it is and, therefore, may increase advertisement in larger individuals without incurring costs to production.

One underexplored hypothesis for the evolution of female ornamentation is their use in female-female competition and social signaling (West-Eberhard 1979; Rosenqvist 1990; Rosvall 2011; Lyon & Montgomerie 2012; Tobias *et al*. 2012). Currently, little is known if the ornament in *S. nigra* is used primarily to signal to males during courtship or in social interactions between females, such as by suppressing display times of rival females. For example, in the sex-role reversed broad snouted pipefish, *Syngnathus typhle*, the visual ornaments displayed on the flanks of females play an important role in female-female competition by functioning as a ‘badge of status’ to intimidate rivals (Berglund & Rosenqvist 2009). If female ornamentation is strongly influenced by social signaling, this may help to maintain honest signaling in the face of negligible costs to fecundity. Future studies could manipulate social context (e.g., interactions between rivals versus potential mates) to elucidate the nature of the female ornament in this species.

Another possible explanation for the origins of the female ornament is that it may have evolved through sensory bias. Studies of sexual selection have shown that mate preference for sexual ornaments can sometimes arise from preexisting perceptual biases (Kirkpatrick & Ryan 1991; Endler 1992; Ryan 1998; Kokko *et al*. 2003; Macías Garcia & Ramirez 2005) that reflect ecological constraints (Endler 1992; Proctor 1992) or basic properties of nervous systems (Ryan & Keddy-Hector 1992; Rosenthal & Evans 1998; Ryan 1998). A classic example of a preexisting female preference for a sexually selected trait is found among fishes in the genus *Xiphophorus* (Basolo 1990; Basolo 1995). Here, female preference for the male ‘sword’ ornament, a colorful extension of the male’s caudal fin, appears to have arisen from a general female preference for larger males (Rosenthal & Evans 1998). In goodeiid fishes, Macías Garcia and Ramirez (2005) demonstrated how such biases, in turn, can evolve into honest sexual signals. Such a possibility in the context of male preferences for body width and stripe pattern of female *S. nigra* warrants further investigation.

To conclude, we found no evidence of a fecundity cost associated with the expression of an extravagant female ornament in a sex role-reversed pipefish, although a potential trade-off between fecundity and egg size was uncovered. Our study adds to a growing body of empirical studies that question whether theoretical assumptions concerning the cost of female ornamentation to fecundity is warranted. Future studies should investigate alternative explanations for the evolution of female ornamentation in this and other species in nature including signaling efficacy, condition dependence, social selection and sensory bias.

## Acknowledgements

We would like to thank Nuno Monteiro for providing helpful comments on an earlier draft.

## Author contributions

MW and BBMW initiated the study. MW performed all dissections with the assistance of DB. KBM, JRM and MW analysed the data. KBM drafted manuscript with help from JRM, MW, DB, BBMW. All authors gave final approval for publication.

